# Breast cancer endocrine therapy exhausts adipocyte progenitors promoting weight gain and glucose intolerance

**DOI:** 10.1101/2020.08.21.259440

**Authors:** Rebecca L. Scalzo, Rebecca M. Foright, Sara E. Hull, Leslie A. Knaub, Stevi Johnson-Murguia, Fotobari Kinanee, Jeffrey Kaplan, Benjamin Freije, Gregory Verzosa, Julie A. Houck, Ginger Johnson, Anni M.Y. Zhang, James D. Johnson, Paul S. MacLean, Jane E.B. Reusch, Sabrina Wright-Hobart, Elizabeth A. Wellberg

## Abstract

Breast cancer survivors treated with anti-estrogen therapies report weight gain and have an elevated risk of type 2 diabetes. Here, we show that current tamoxifen use associated with larger breast adipocyte diameter only in women with a BMI >30 kg/m^2^. To understand the mechanisms behind these clinical findings, we investigated the impact of estrogen deprivation and tamoxifen in a relevant pre-clinical model of obesity. Specifically, mature female mice were housed at thermoneutrality and fed either a low-fat/low-sucrose (LFLS) or a high-fat/high-sucrose (HFHS) diet. Consistent with the high expression of *Esr1* observed in single-cell RNA sequencing of mesenchymal stem cells from adipose tissue, endocrine therapies induced adipose accumulation and preadipocyte expansion, but resulted in adipocyte progenitor depletion only in the context of HFHS. Consequently, 7-week endocrine therapy supported adipocyte hypertrophy and was associated with hepatic steatosis, hyperinsulinemia, insulin resistance, and glucose intolerance, particularly in HFHS fed females. Metformin or pioglitazone, glucose lowering drugs used to treat diabetes, prevented the effects of tamoxifen but not estrogen deprivation on adipocyte size and insulin resistance in HFHS-fed mice. This translational study suggests that endocrine therapies act via ERα to directly disrupt adipocyte progenitors and support adipocyte hypertrophy, leading to ectopic lipid deposition that may promote hyperinsulinemia, insulin resistance and type 2 diabetes. Interventions that target insulin action should be considered for some women receiving life-saving endocrine therapies for breast cancer.

## Introduction

Breast cancer is the second most common cause of death among women, and >70% of breast cancer patients are diagnosed with estrogen receptor (ER) positive tumors ^1^. In premenopausal women, ER-positive cancers are often treated using selective ER modulators (SERMs) ^2^, the most common of which, tamoxifen (TAM), is prescribed at the time of diagnosis and is used for 5-10 years (until biological menopause) ^3^. After menopause or during ovarian function suppression, aromatase inhibitors are used to block peripheral estrogen production ^4^. SERMs and aromatase inhibitors significantly reduce the risk of breast cancer recurrence and have saved millions of lives ^5^. Notwithstanding the importance of ER-targeted therapies for breast cancer, use of these agents is linked to type 2 diabetes ^6-9^. A recent report suggested that up to 48% of diabetes cases in breast cancer survivors are attributable to the effects of endocrine therapy ^10^, and adverse metabolic effects such as hepatic steatosis and diabetes appear to be more prevalent in women with overweight or obesity ^11,12^. Type 2 diabetes is a leading cause of morbidity and premature mortality and is a risk factor for breast cancer recurrence ^13,14^.

ER is targeted in breast cancer because of its mitogenic role; however, estrogen maintains energy expenditure and adipose tissue homeostasis in premenopausal women ^15^. Activation of ER decreases energy intake and increases energy expenditure; together these responses to ER activation contribute to the maintenance of body mass over time. Further, ER activation attenuates visceral adipose storage and supports “healthy” adipose tissue expansion in subcutaneous depots through preadipocyte recruitment ^16,17^. Disrupted ER signaling within adipose tissue may interfere with normal adipocyte biology and contribute to the elevated risk for diabetes, or potentially to weight gain, in women treated with endocrine therapy. Conflicting metabolic consequences of endocrine therapy have been reported in preclinical models. A number of metabolic studies employed supra-physiological doses of TAM to induce the expression of Cre-ER transgenes (reviewed in ^18^). High-dose TAM over a short period of time leads to weight loss in mice, but this is not representative of the physiological response in breast cancer survivors ^19-22^. Importantly, few reports have considered the interaction with excess adiposity, which impacts over 70% of adult women ^23^. The rodent studies where TAM was administered at a clinically relevant dose focus on anti-cancer effects and have not reported whole-body metabolic outcomes ^24,25^.

To gain insight into the mechanisms linking endocrine therapy to diabetes, insulin resistance and adiposity, we conducted an analysis of breast adipose tissue biopsies from tamoxifen-treated women and investigated the impact of breast cancer anti-estrogen treatment in mature female mice housed at thermoneutrality and fed either a low-fat/low-sucrose (LFLS) or a high-fat/high-sucrose (HFHS) diet. Human breast adipose tissue from women taking TAM exhibited significantly increased adipocyte diameter only when BMI was > 30 kg/m^2^. In a mouse model, we found that long-term TAM treatment increased fat gain in HFHS-fed but had no effect on body mass or fat in LFLS-fed female mice. In contrast, estrogen deprivation promoted fat gain regardless of diet and adiposity. In the context of a HFHS diet, endocrine therapies inhibited expansion of adipose progenitor cells (Lin^-^ /CD29^+^/CD34^+^/Sca1^+^/CD24^+^), promoted adipocyte hypertrophy, and supported glucose intolerance and ectopic lipid deposition. In the rodent model, the glucose lowering drugs metformin and pioglitazone reduced adipocyte hypertrophy and restored insulin sensitivity in the presence of TAM but not during estrogen deprivation. Understanding the metabolic impact of endocrine therapy is important for preventing diabetes as well as breast cancer-specific adverse outcomes, such as metastasis- or progression-free survival. Overall, our study demonstrates metabolic consequences of breast cancer endocrine therapy that are reported in patients and highlights the need to consider adjunctive interventions to combat the negative metabolic effects of endocrine therapies in breast cancer survivors.

## Results

### TAM exposure is associated with adipocyte hypertrophy in women

Links between endocrine therapy use and diabetes as well as hepatic steatosis have been reported ^5-7,10,11^; however, the impact of these treatments on weight gain are controversial and there are few studies on adipose tissue effects in women. Based on this evidence, we analyzed human breast adipose tissue and BMI data from women who were currently taking prescribed tamoxifen and compared that to women who were not taking endocrine therapy. We found that the average adipocyte diameter significantly correlated with BMI (Figure 1A; P = 0.0001), validating what we previously found ^23^. When we analyzed adipocytes from women classified as lean (BMI < 25 kg/m^2^), overweight (BMI 25-29 kg/m^2^) or obese (BMI > 30 kg/m^2^) we found that current tamoxifen use associated with larger adipocyte diameter only in those women with obesity (Figure 1B-C; P = 0.03), compared to women classified as lean or overweight (Figure 1B-C). For those with BMI > 30 kg/m^2^, the distribution of adipocyte sizes in tamoxifen users reflected a greater proportion of large cells compared to women not taking tamoxifen (Figure 1D). These data suggest that endocrine therapy, particularly tamoxifen, impacts adipose tissue, which may relate to the elevated type 2 diabetes risk observed in breast cancer survivors.

**Figure 1.**
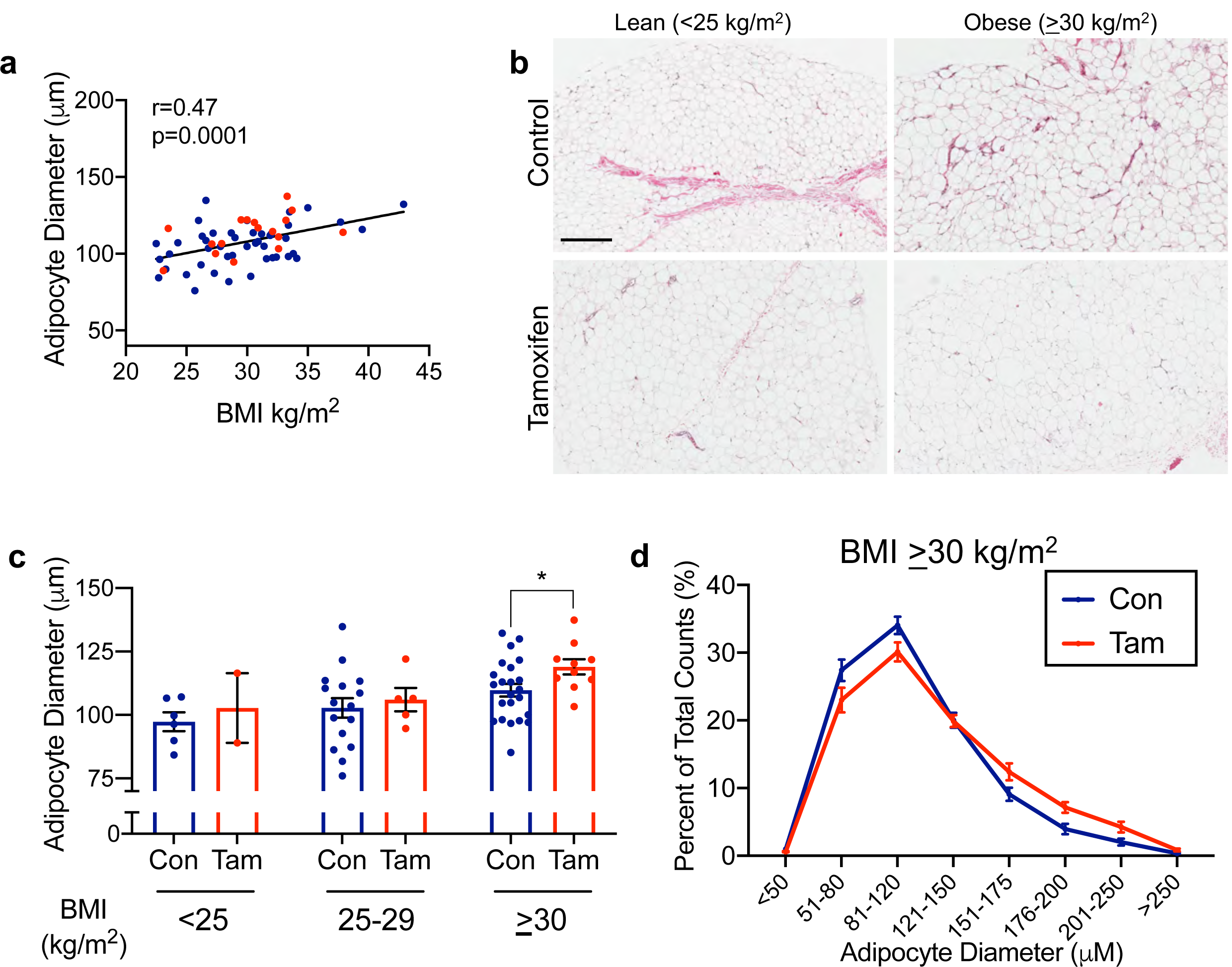
Tamoxifen treatment associates with larger adipocytes in women with BMI > 30 kg/m^2^. **a**, spearman correlation of subject BMI and average adipocyte diameter. Red circles are from tamoxifen-treated women, blue circles are controls. **b**, representative breast adipose images from one subject each, control or tamoxifen-treated, BMI < 25 kg/m^2^ or > 30 kg/m^2^. Scale bar is 400 µm. **c**, average adipocyte diameters from control or tamoxifen-treated women with BMI <25 kg/m^2^ (Con n=6, Tam n=2), 25-29 kg/m^2^ (Con n=16, Tam n=5), or >30 kg/m^2^ (Con n=22, Tam n=10). p<0.05 by unpaired t-test. **d**, Adipocyte diameter represented as the proportion of each cell size representative to total cell count in women with BMI > 30kg/m^2^. Note the frequency of larger adipocytes in tamoxifen-treated women. Con n=22, Tam n=10.

### Modeling effects of TAM and EWD in female mice

The significant changes in adipocyte size seen in women taking tamoxifen reveal questions that can only be studied in pre-clinical animal models. To determine the effect of excess adiposity (i.e. obesity) on the metabolic response to endocrine therapy, we fed either low fat/low sucrose (LFLS) or high fat/high sucrose (HFHS) diets to female mice at thermoneutrality ^26^ (∼30C) to promote fat gain (Figure 2A). Mature, ovariectomized mice were given supplemental estradiol (E_2_) and then randomized to endocrine treatments based on body fat percentage within diet groups (Figure 2A). Mice were maintained on E_2_ or given TAM in the presence of E_2_ (E_2_+TAM), or supplemental E_2_ was withdrawn (EWD), as would occur with aromatase inhibition. We previously showed that adipose tissue from rats ^27^ and mice ^26^ does not express detectable levels of the *Cyp19a1* gene, which encodes the aromatase enzyme. This was verified in a recently published compendium of single-cell sequencing data from mouse adipose and mammary tissue (Tabula Muris) ^28^. The effectiveness of the hormonal manipulations in female mice was confirmed by measuring uterine mass after 2 weeks of treatment (Figure 2B). Compared to control E_2_-supplemented mice, uterine masses were lower in the presence of E_2_+TAM (Figure 2B-C; *P* = 0.0005) and after EWD (Figure 2B-C; *P* < 0.0001). Thus, our pre-clinical mouse model is sufficient for determining the effects estrogen withdrawal and tamoxifen in a relevant physiological setting.

**Figure 2.**
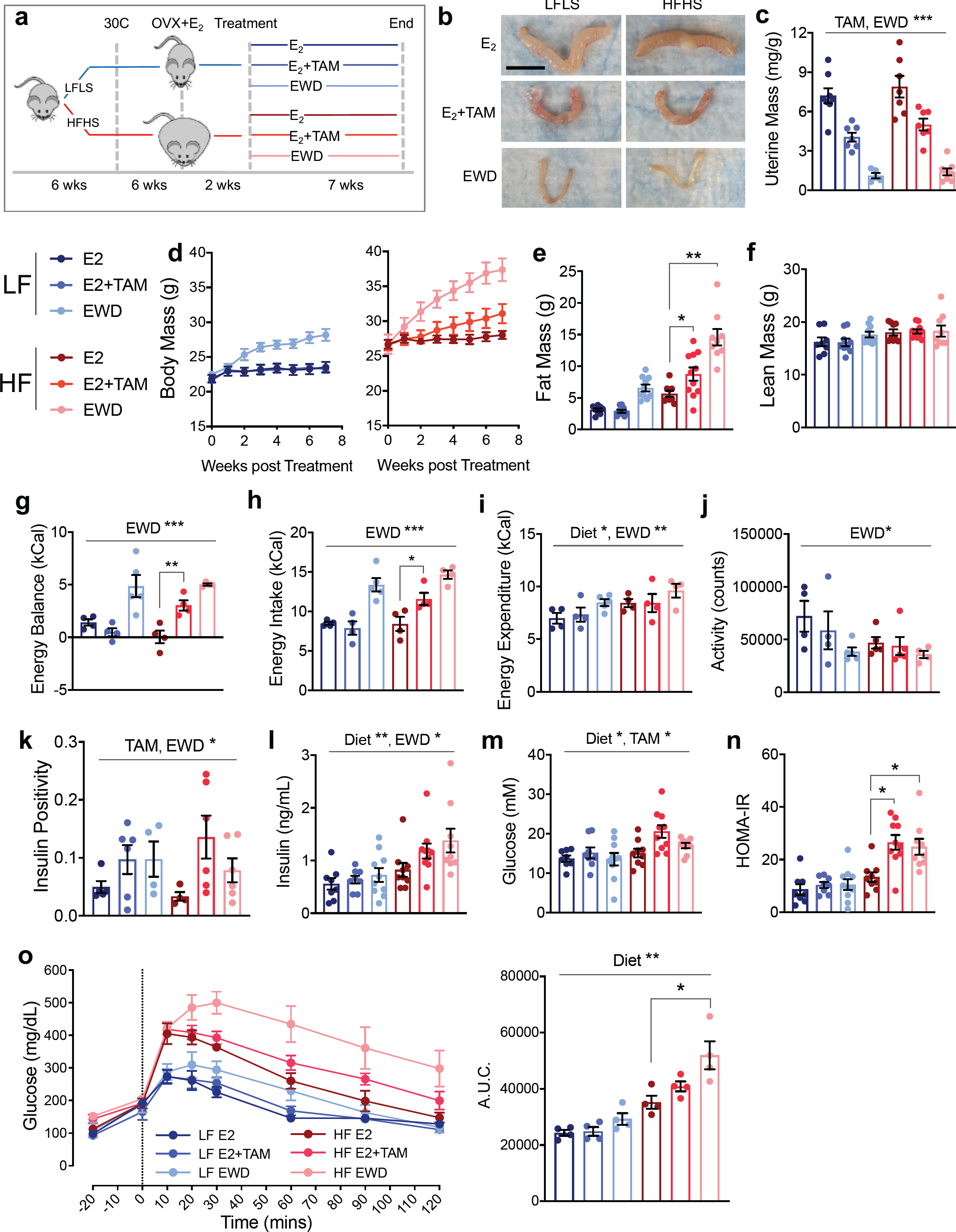
Endocrine therapy promotes weight gain and glucose intolerance in HFHS fed females. **a**, juvenile wild type C57Bl/6 mice are fed a low fat/low sucrose (LFLS, blue lines) or high fat/high sucrose (HFHS, red lines) diet and housed at ∼30C. Mature females are ovariectomized (OVX) and supplemented with estradiol (E_2_) for 2 weeks. Mice are then randomized to one of three treatments within each diet group: E_2_ maintenance (E_2_), E_2_ plus tamoxifen (E_2_+TAM) or withdrawal of supplemental E_2_ (EWD). Treatments continue for 7 weeks. **b**, representative images of uteri from one mouse in each diet/treatment group. Scale bar is 1 cm. **c**, uterine mass presented as mg/g body mass for each group: LFLS E_2_ (n=8), LFLS E_2_+TAM (n=7), LFLS EWD (n=5), HFHS E_2_ (n=7), HFHS E_2_+TAM (n=7), HFHS EWD (n=8). Main effects of TAM and EWD treatments (P ≤ 0.001) by 2-way ANOVA. **d**, Body mass (grams) of mice in each treatment group beginning at the start of treatment and continuing for 7 weeks: LFLS E_2_ (n=8), LFLS E_2_+TAM (n=9), LFLS EWD (n=10), HFHS E_2_ (n=9), HFHS E_2_+TAM (n=12), HFHS EWD (n=9). For LFLS mice, EWD resulted in greater weight gain over time (P < 0.0001 for time x EWD interaction) but E_2_+TAM did not, compared with E_2_ alone. For HFHS mice body weight gain was greater over time with both TAM and EWD (P = 0.001 and P < 0.0001 respectively). 2 × 3 ANOVA (factors: time, diet, and treatment) with interaction tests determined significance. **e**, fat mass in grams for each group LFLS E_2_ (n=8), LFLS E_2_+TAM (n=9), LFLS EWD (n=10), HFHS E_2_ (n=9), HFHS E_2_+TAM (n=11), HFHS EWD (n=9). Fat mass was greater in HFHS mice with TAM and EWD (P = 0.02 and P = 0.001 respectively; interactions by 2-way ANOVA (factors: diet and individual treatment). **f**, lean body mass in grams for each group. Sample sizes are the same as for fat mass. **g**, 24-hour average energy balance in kCal. (P < 0.001 for the main effect of EWD; P < 0.01 for the interaction). **h**, 24-hour average energy intake in kCal. (P < 0.001 for the main effect of EWD; P < 0.05 for the interaction). **i**, 24-hour average energy expenditure in kCal. (P < 0.05 and P < 0.01 respectively for the main effects). **j**, 24-hour average physical activity counts. (P < 0.05). **k**, the proportion of insulin positive out of total pixels. (p<0.05; factors: diet and individual treatments; 2×2 ANOVA); **l** and **m**, serum insulin and glucose were measured in samples taken at sacrifice following a 4 hour fast. LFLS E_2_ (n=8), LFLS E_2_+TAM (n=9), LFLS EWD (n=10), HFHS E_2_ (n=9), HFHS E_2_+TAM (n=11), HFHS EWD (n=9). Effect of diet insulin and glucose P < 0.05. Effect of EWD on insulin P = 0.03. Effect of TAM on glucose P = 0.02. **n**, The Homeostatic Model Assessment of Insulin Resistance (HOMA-IR) was calculated from fasting insulin and glucose. Both EWD and TAM administration resulted in greater insulin resistance in the context of HFHS [P = 0.04 and P = 0.01 respectively; significance represents the statistical interactions from 2×2 ANOVA (factors: diet and individual treatments)]. **o**, Oral glucose tolerance tests were performed, and the glucose excursion is presented as the area under the curve. Both EWD and TAM exacerbated the worsening of glucose tolerance observed with HFHS (P = 0.04 and P = 0.06 respectively for the statistical interaction). LFLS E_2_ (n=8), LFLS E_2_+TAM (n=9), LFLS EWD (n=10), HFHS E_2_ (n=9), HFHS E_2_+TAM (n=11), HFHS EWD (n=9).

### Tamoxifen and EWD promote fat gain and impair glucose tolerance

Suppression of ovarian function, which occurs during menopause or with ovariectomy, associates with adipose accumulation and redistribution in women and mice ^29,30^. While weight gain is a potential and concerning anecdotal side effect of breast cancer therapy, clinical studies report inconsistent data due to differences in study design, group comparisons, and analysis timelines ^31^. We evaluated body mass and composition in female mice fed LFLS or HFHS diets. Overall, EWD treated mice gained more weight compared with E_2_-treated mice in each diet group (*P* < 0.001). In contrast to several published studies ^19-22,32^, TAM treatment promoted rapid weight gain, but only in HFHS fed mice (*P* = 0.001). Evaluation of body composition showed significant interactions for diet and TAM (*P* = 0.02), as well as diet and EWD (*P* = 0.001) for fat mass (Figure 2B) but not for lean mass (Figure 2C).

To determine the physiological mechanisms driving excess weight gain, whole-animal calorimetry was performed during the second week of endocrine therapy. This time frame marked the beginning of the body weight separation seen in both LFLS and HFHS mice and represents a dynamic phase where differences in energy intake and expenditure could be appreciated. TAM treatment created a greater positive energy balance in the HFHS mice compared with the LFLS mice (Figure 2G; *P* = 0.007), while both diet groups experienced a positive energy balance during EWD (Figure 2G; P < 0.0001). In all cases, the positive energy balance was associated with greater energy intake (Figure 2H; EWD *P* < 0.0001; TAM *P* = 0.017). Greater energy expenditure was seen in the HFHS mice compared with LFLS mice (Figure 2I; P < 0.02) and during EWD regardless of diet (P = 0.004). Physical activity counts tended to be lower in HFHS compared to LFLS mice (Figure 2J; P = 0.06) and were significantly lower during EWD (P = 0.012) but were unaffected by TAM treatment.

The most consistently reported adverse metabolic effect of breast cancer endocrine therapy is an elevated risk of type 2 diabetes ^6-8,18^. Therefore, we evaluated fasting insulin, glucose, and calculated HOMA-IR, an estimation of insulin resistance, as early markers that could predict diabetes. By seven weeks, insulin was elevated in HFHS versus LFLS mice (Figure 2K; P < 0.01) and was higher during EWD (P = 0.029); however, the effect of TAM was not significant (P = 0.09). Increased fasting insulin can be a product of increased physical beta-cell mass and/or relative insulin hypersecretion. We estimated beta-cell mass by quantifying insulin-positive area in pancreas sections ^33^. With both TAM and EWD treatments, beta-cell area was greater regardless of diet (Supp Fig 1A-B; P < 0.05). However, at this time point only the HFHS mice showed elevated circulating insulin levels (Figure 2K). These data indicate that, while blocking ER signaling is associated with increased physical beta-cell mass in this model, elevated circulating insulin levels are only seen in the context of HFHS diet.

Glucose was also elevated in HFHS versus LFLS mice (Figure 2L; P < 0.05) and was elevated in both diet groups by TAM (P = 0.02), but not EWD. The distinct effects of endocrine therapies on insulin and glucose resulted in elevated HOMA-IR in the HFHS TAM-(P = 0.01) and EWD-treated (P = 0.04) mice (Figure 2M). We also found that glucose tolerance was poorer overall in HFHS mice (greater glucose area under the curve) during an oral glucose tolerance test (Figure 2N-M; *P* < 0.001). Within this diet group, glucose tolerance tended to be worse with TAM treatment (*P* = 0.06) and was significantly impaired during EWD (*P* = 0.04). Despite the observation that fasting insulin and glucose were greater with TAM, particularly in HFHS mice, it did not appear to impair glucose tolerance at the same time point. Overall, these data suggest that endocrine therapies adversely affect glucose tolerance and insulin sensitivity in HFHS fed females. Moreover, data from the LFLS mice indicate that fat gain, which was observed with EWD regardless of diet, is not always associated with impaired glucose tolerance.

### Estrogen receptor is expressed in progenitor cells from metabolic tissues

To gain further understanding of what cell types respond to changes in estrogen and *Esr1* (ERα) signaling, we explored the Tabula Muris resource of single cell sequencing data from numerous murine tissues (without uterus, ovaries, and testes) ^28^. Interestingly, the highest expression levels of *Esr1* were detected in cells from adipose (fat; stromal fraction), skeletal muscle, mammary tissue, and liver (Figure 3A). Within adipose tissue, *Esr1* was most highly expressed in a small population of unannotated, myeloid-derived cells that express CD45 (*Ptprc*), CD29 (*Itgb1*), Sca1 (*Ly6a*), and adipocyte progenitor markers *Dpp4* and *Pi16*^34^, but lack *Pdgfra*, CD11b (*Itgam*), and Cd34 (Figure 3A-B)^28^. These cells are similar to those previously described as a bone-marrow derived population of adipose progenitor cells ^35-37^. *Esr1* was also highly expressed in mesenchymal stem cells from both adipose tissue (Figure 3B) and skeletal muscle (Figure 3C). Within skeletal muscle, the satellite cells (Figure 3C), a self-renewing muscle cell that contributes to tissue repair ^38^, also highly expressed *Esr1*. In the mammary tissue, stromal cells and a subset of luminal epithelial cells demonstrated high *Esr1* levels (Figure 3D). The expression pattern of *Esr1* in adipose, mammary stroma, muscle, and liver suggests that breast cancer endocrine therapy may target *Esr1* in progenitor cell populations, particularly in tissues that impact whole-body metabolism.

**Figure 3.**
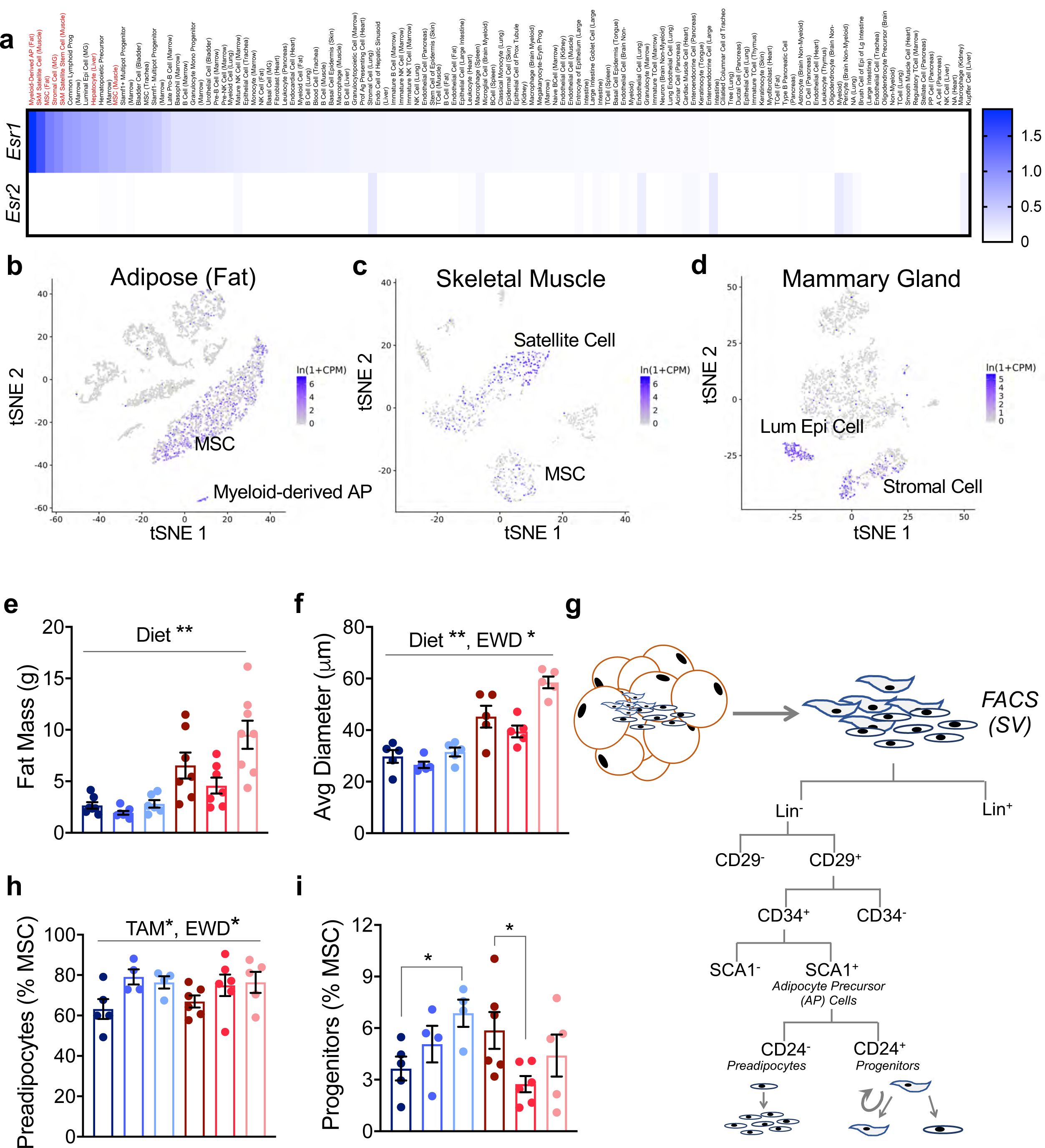
Tamoxifen and EWD deplete adipocyte progenitors after 2 weeks of treatment. **a**, heatmap of *Esr1* and *Esr2* gene expression from the Tabula Muris single cell RNA sequencing resource. The cell type and tissue are listed. In red are *i)* myeloid-derived adipose progenitor; *ii)* skeletal muscle satellite cell; *iii)* mesenchymal stem cell (MSC) from adipose; *iv)* stromal cell from mammary gland; *v)* skeletal muscle satellite stem cell; *vi)* hepatocyte; and *vii)* MSC from skeletal muscle. **b-d**, tSNE plots of *Esr1* expression in FACS separated cells from adipose (b), skeletal muscle (c), and mammary gland (d) downloaded from the Tabula Muris website. **e**, Fat mass in grams of mice in each group after 2 weeks of treatment: LFLS E_2_ (n=7), LFLS E_2_+TAM (n=7), LFLS EWD (n=6), HFHS E_2_ (n=7), HFHS E_2_+TAM (n=7), HFHS EWD (n=8). Fat mass was greater in HFHS mice compared with LFLS (diet effect, P = 0.01). **f**, Average adipocyte diameter measured histologically after 2 weeks of treatment: n=5 per group. Average adipocyte diameter was greater in HFHS mice and EWD mice (P < 0.0001 and P = 0.02 respectively). **g**, FACS strategy to evaluate the proportion of adipose mesenchymal stem cells that were composed of preadipocytes and progenitor cells after 2 weeks of treatment. **h**, Proportion of adipose mesenchymal stem cells that were composed of preadipocytes. TAM and EWD treatment resulted in a greater proportion of preadipocyte cells irrespective of diet (P < 0.05 for each main effect). **i**, Proportion of adipose mesenchymal stem cells that were composed of progenitor cells. TAM treatment resulted in a smaller proportion of progenitor cells in HFHS mice (P = 0.02) while with EWD, the proportion of progenitor cells was greater in LFLS mice and unchanged in HFHS mice (P = 0.04). Significance represents the statistical interactions from 2×2 ANOVA (factors: diet and individual treatments). LFLS E_2_ (n=5), LFLS E_2_+TAM (n=4), LFLS EWD (n=4), HFHS E_2_ (n=6), HFHS E_2_+TAM (n=6), HFHS EWD (n=5) for h and i.

### TAM and EWD differentially impact adipose tissue expansion in LFLS and HFHS mice

Our previous study in a model of obesity and ER-positive breast cancer suggested a role for expanding subcutaneous adipose tissue and adipocyte hypertrophy in endocrine therapy resistance after menopause ^26^. Based on this work, and on our observations of predominant *Esr1* expression in progenitor cells from adipose tissue, we evaluated adipocyte precursor cell types in the subcutaneous adipose from a cohort of mice after two weeks of treatment ^39^. At this time point, HFHS had greater fat mass than LFLS females, but there were no effects of endocrine therapy (Figure 3E). The average adipocyte diameter (Figure 3F) was larger overall in HFHS fed mice (P ≤ 0.0001) and during EWD treatment (P = 0.02). Using FACS ^40^, we evaluated the proportion of adipose mesenchymal stem cells that were composed of preadipocytes and progenitor cells (Figure 3G-I; Supplemental Figure 2). The fraction of preadipocytes in both diet groups was greater with TAM and EWD treatments (Figure 3H; *P* = 0.017). In the progenitor population (Figure 3I), we found a significant interaction between diet and both TAM (P = 0.02) and EWD (P = 0.04) treatments, where progenitors were greater in the LFLS mice, but lower (TAM) or unchanged (EWD) in the HFHS mice. Regardless of diet, disrupting ER signaling appears to stimulate an early burst in preadipocyte proliferation; however, formation of new cells may not be sustainable in HFHS mice, as the source of these preadipocytes (progenitor cells) is concomitantly being depleted.

Next we investigated the subcutaneous adipose after seven weeks of treatment to evaluate adipose tissue expansion in response to endocrine therapy. Subcutaneous adipose tissue mass was greater with both TAM (P = 0.002) and EWD (P = 0.01) treatments in HFHS mice (Figure 4A-B). We also saw a slightly more frequent occurrence of larger adipocytes in LFLS females after EWD (Figure 4B-C). The greatest proportion of large adipocytes was seen in TAM and EWD treated HFHS females (Figure 4B-C). The average diameter of adipocytes in this depot was significantly greater in HFHS mice by TAM (P = 0.04) and EWD (P = 0.01) treatments (Figure 4D). When combined with the potential depletion of adipocyte progenitors observed two weeks after treatment, the adipocyte hypertrophy seen in HFHS mice after longer-term (seven week) endocrine therapy is consistent with a depletion of progenitor cells and inability to expand adipose tissue by hyperplasia. Notably, we observed similar adipocyte hypertrophy after TAM treatment in women with an elevated BMI (Figure 1).

**Figure 4.**
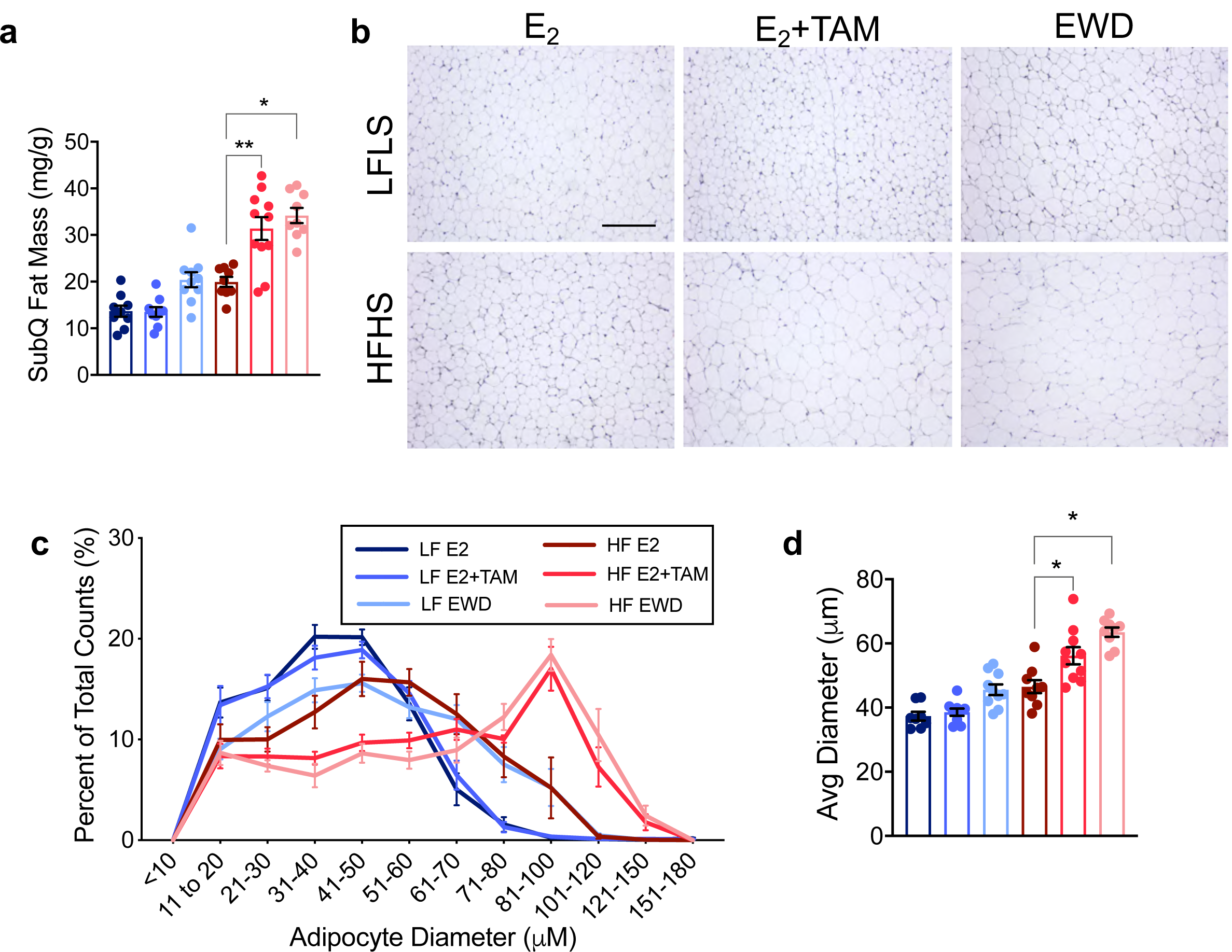
Adipocyte hypertrophy after 7 weeks of endocrine therapy. **a**, Subcutaneous fat mass presented as mg/g in mice from each treatment group after 7 weeks of TAM or EWD: LFLS E_2_ (n=9), LFLS E_2_+TAM (n=9), LFLS EWD (n=10), HFHS E_2_ (n=9), HFHS E_2_+TAM (n=11), HFHS EWD (n=9). There was greater accumulation of fat mass in HFHS mice with both TAM and EWD (P = 0.002 and P = 0.01 respectively. Significance represents the statistical interactions from 2×2 ANOVA (factors: diet and individual treatments). **b**, Representative images of adipocytes from one mouse in each diet/treatment group. Scale bar is 500 µm. **c**, Adipocyte diameter represented as the proportion of each cell size representative to total cell count. Note the frequency of larger adipocytes in HFHS females after EWD and TAM treatment. LFLS E_2_ (n=8), LFLS E_2_+TAM (n=9), LFLS EWD (n=10), HFHS E_2_ (n=9), HFHS E_2_+TAM (n=11), HFHS EWD (n=9). **d**, Quantification of adipocyte size represented by the mean adipocyte diameter. The average diameter of adipocytes in this depot was greater in HFHS mice treated with TAM and EWD (P = 0.04 and P = 0.01 respectively). Significance represents the statistical interactions from 2×2 ANOVA (factors: diet and individual treatments). Sample sizes are the same as in c.

### Increased hepatic steatosis and altered hepatic metabolism with E_2_ withdraw

Several clinical studies report hepatic steatosis in breast cancer patients after treatment with TAM or aromatase inhibitors ^41-43^. Adipocyte hypertrophy and decreased capacity for adipose fuel storage is known to associate with lipid spill-over into circulation and to present with ectopic lipid in other tissues, including the liver, in both humans and animal models ^44,45^. We found that EWD treatment worsened steatosis compared to E_2_ in HFHS and LFLS mice (Figure 5A; P < 0.05) without affecting liver mass (Figure 5B). While HFHS TAM-treated mice demonstrated hepatic steatosis, there was not a significant effect of TAM treatment on this outcome (Figure 5A). Given our observation that ectopic fat accumulated in the liver, we also examined pancreatic fat accumulation, which has also been linked to beta-cell function ^33^. Estimates of pancreatic fat content were variable and not significantly different between any groups (Supplemental Figure 1C). Collectively, these observations indicate that endocrine therapy can affect ectopic lipid deposition, but there may be more prominent direct effects on cells that express *Esr1* within susceptible tissues such as the liver.

**Figure 5.**
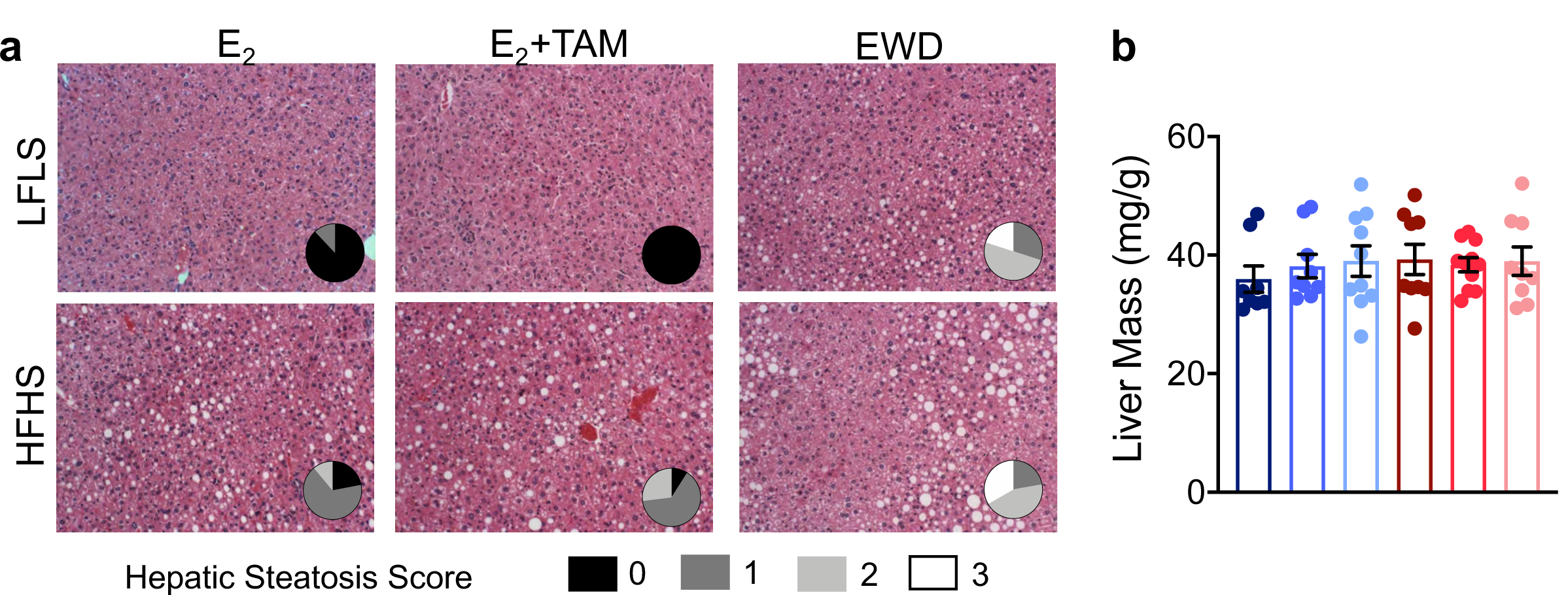
Endocrine therapy associates with hepatic steatosis. **a**, H&E stained sections of livers from one mouse in each group. Pie chart insets represent quantification of hepatic steatosis severity in each group. Score legend is shown below the H&E images. LFLS E_2_ (n=8), LFLS E_2_+TAM (n=9), LFLS EWD (n=10), HFHS E_2_ (n=9), HFHS E_2_+TAM (n=11), HFHS EWD (n=9). **b**, liver mass presented as grams per gram body mass. LFLS E_2_ (n=8), LFLS E_2_+TAM (n=9), LFLS EWD (n=10), HFHS E_2_ (n=9), HFHS E_2_+TAM (n=11), HFHS EWD (n=9).

In rodent models of diet-induced obesity, hepatic lipid oxidation (mitochondrial function) is lower and hepatic lipid storage is greater ^46,47^. To determine the impact of EWD and TAM administration on hepatic mitochondria, we evaluated mitochondrial respiration and content. Mitochondrial respiration was assessed with both a carbohydrate-linked SUIT (PMGS) and a lipid-linked SUIT (PCMS). The ratio of leak to ADP stimulated respiration (L/P ratio) with PMGS was greater in the HFHS mice with EWD compared with the HFHS E_2_ mice and was lower in the LFLS EWD mice compared with LFLS E_2_ mice (Supplemental Fig 3B; *P* = 0.04). TAM administration did not affect this measure. There was no effect of diet or either endocrine therapy on the ratios measured in the PCMS suit (Supplemental Fig 3D-F). The only significant effect of diet or either treatment on mitochondrial complex content was seen in complex III, which was lower with EWD in both LFLS and HFHS mice (Supp Fig 3G-H; P =0.012). For all other complexes, neither TAM nor EWD treatment impacted protein content in either diet group (Supplemental Fig 3G-H). These data suggest that the liver phenotype observed in HFHS mice treated with TAM or EWD is not the product of direct action of TAM or EWD on hepatic mitochondria after 7 weeks of treatment; however, disruption of ER signaling is known to directly impact the liver in several studies^48,49^.

### Metformin and pioglitazone improve measures of metabolic dysfunction during TAM but not EWD

The impact of endocrine therapy on adipose tissue expansion, glucose tolerance, and insulin resistance suggested the potential for anti-hyperglycemic therapies to prevent adverse metabolic effects of breast cancer therapies. We administered either metformin or pioglitazone to HFHS females treated with TAM or EWD to determine whether either intervention could lower HOMA-IR and/or reduce adipocyte hypertrophy compared to mice treated with either endocrine therapy alone. Metformin primarily acts by lowering hepatic glucose output, while pioglitazone activates PPAR isoforms to augment insulin sensitivity. In adipose, pioglitazone promotes adipocyte differentiation; in liver pioglitazone can decrease hepatic lipid content ^50-52^.

Total fat mass (Figure 6A) in TAM treated mice was unaffected by metformin but was lower with pioglitazone (P = 0.05). Neither intervention altered total fat mass during EWD (Figure 6A). Despite no significant effects on adipose mass, average adipocyte diameter was smaller with both metformin (P = 0.01) and pioglitazone (P = 0.001) in TAM treated mice (Figure 6B). A similar effect was seen with pioglitazone treatment during EWD (P = 0.003), but not with metformin treatment. The distribution of adipocyte sizes coordinately showed a greater proportion of small adipocytes with both metformin and pioglitazone treatment of TAM females, and only with pioglitazone treatment during EWD (Figure 6C). Overall, these data indicate that anti-hyperglycemic drugs may directly or indirectly stimulate adipocyte hyperplasia to reduce overall cell size without dramatically reducing tissue mass during TAM and EWD treatments.

**Figure 6.**
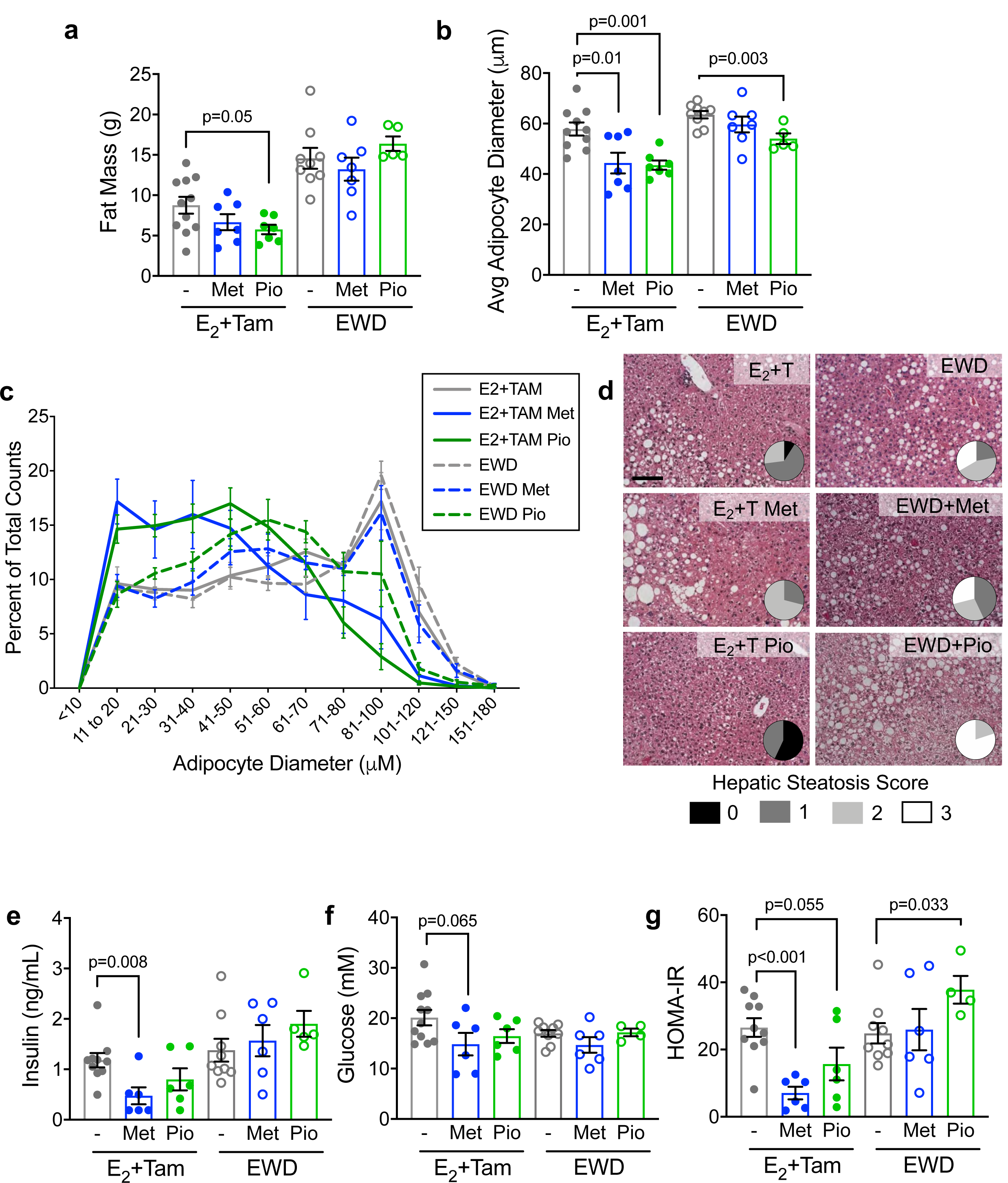
Anti-hyperglycemic drugs improve metabolic outcomes during TAM treatment. **a**, Fat mass in grams from TAM or EWD treated females that received no interventions (-), metformin (Met), or pioglitazone (Pio) for 7 weeks. Pio treatment associated with reduced fat mass during TAM treatment (p=0.05 by unpaired t-test). E_2_+TAM no intervention (n=11), Met (n=7), Pio (n=7); EWD no intervention (n=9), Met (n=7), Pio (n=5). **b**, Average adipocyte diameter from subcutaneous adipose tissue. In TAM treated females, both Met (p=0.01) and Pio (p=0.001) associated with smaller diameter by unpaired t-test. In EWD treated females, only Pio associated with smaller diameter (p=0.003 by unpaired t-test). E_2_+TAM no intervention (n=10), Met (n=7), Pio (n=7); EWD no intervention (n=9), Met (n=7), Pio (n=5). **c**, Adipocyte diameter represented as the proportion of each cell size representative to total cell count. Sample sizes are the same as in b. **d**, H&E stained liver sections from one mouse in each group. Pie chart insets represent quantification of hepatic steatosis severity in each group. Score legend is shown below the H&E images. E_2_+TAM no intervention (n=11), Met (n=7), Pio (n=7); EWD no intervention (n=9), Met (n=7), Pio (n=5). **e** and **f**, serum insulin and glucose were measured in samples taken at sacrifice following a 4 hour fast. E_2_+TAM no intervention (n=11), Met (n=7), Pio (n=7); EWD no intervention (n=9), Met (n=7), Pio (n=5). In only E_2_+TAM mice, insulin and glucose were lower with Met (P < 0.01 and P = 0.065 respectively). **g**, The Homeostatic Model Assessment of Insulin Resistance (HOMA-IR) was calculated from fasting insulin and glucose measures as a surrogate measure of insulin resistance. In E_2_+TAM mice, both Met and Pio treatment lowered HOMA-IR (P < 0.001 and P = 0.055 respectively). In EWD mice, administration of Pio resulted in greater HOMA-IR (P = 0.033).

### Differential effects of metformin and pioglitazone on hepatic steatosis in TAM and EWD females

We postulated that the generation of new adipocytes would provide sites for storing excess circulating substrates and prevent ectopic lipid deposition and evaluated whether either intervention improved hepatic steatosis during endocrine therapy in HFHS females. Surprisingly, during TAM treatment, metformin tended to worsen (Figure 6D; p=0.06), while pioglitazone significantly improved hepatic steatosis (Figure 6D; p=0.02). There were no effects of either intervention on liver mitochondrial respiration in the context of TAM treatment (Supplemental Figure 4A-F). Liver mitochondrial complex I was lower with metformin in TAM treated females (Supplemental Figure 4G-H; p=0.04), but neither intervention impacted liver mitochondrial content regardless of endocrine treatment. No effect of metformin was seen on hepatic steatosis during EWD (Figure 6D), and in contrast to TAM treatment, pioglitazone tended to worsen this outcome (Figure 6D; p=0.08). During EWD, we found that with pioglitazone PM L/P (Supplemental Figure 4A; p=0.069) and PMGS L/P (Supplemental Figure 4B; p=0.078) tended to be lower, but there were no other effects of metformin or pioglitazone on mitochondrial respiration. Overall, these data suggest that anti-hyperglycemic drugs may differentially impact hepatic steatosis depending on the presence or absence of TAM, but these effects do not appear to be the consequence of an altered hepatic mitochondrial phenotype.

### Improved insulin sensitivity with metformin during TAM treatment but not EWD

Metformin and pioglitazone are widely prescribed to lower circulating insulin and glucose levels in patients with type 2 diabetes; therefore, we evaluated these parameters in mice. While metformin had no effect on overall adipose tissue mass (Figure 6A) and hepatic steatosis was worse with its administration (Figure 6D), circulating insulin was lower with metformin in the presence of TAM (Figure 6E; p=0.008) and glucose tended to be as well (Figure 6F; p=0.065). Together these effects resulted in lower HOMA-IR (Figure 6G; p<0.001). Conversely, insulin (Figure 6E) and glucose (Figure 6F) were modestly less with pioglitazone, leading to a non-significant lowering of HOMA-IR (Figure 6G; p=0.055) during TAM treatment. In contrast, during EWD treatment metformin had no effect on insulin, glucose, or HOMA-IR (Figure 6E-G). Although pioglitazone did not significantly impact insulin or glucose independently, when the measures were combined, HOMA-IR was greater during EWD (Figure 6G; p=0.033). Together, these data suggest that each anti-hyperglycemic drug uniquely impacts the adipose, liver, and circulating insulin and glucose. Importantly, these effects were distinct depending on whether TAM or E_2_ were present, which has implications for women receiving different classes of endocrine therapies.

## Discussion

Here we report, to our knowledge, the first preclinical study to investigate the metabolic changes that occur in obese females taking widely-used breast cancer endocrine therapies, and the first evidence that TAM may affect adipose tissue in women. We found that breast cancer endocrine therapies ^24-26^ promote similar metabolic derangements in HFHS fed mice that are reported in breast cancer patients ^7,8,12,42,53-57^, including weight and fat gain, greater energy intake, elevated fasting insulin and glucose, adipocyte hypertrophy, and hepatic steatosis (summarized in Supplemental Table 1). The increase in hepatic steatosis and type 2 diabetes in breast cancer patients ^7,8,12,42,53-55^ is thought to be more common in women with overweight or obesity ^11,12^. Consistent with this model, we observed that TAM use associated with adipocyte hypertrophy in women with a BMI >30kg/m^2^. While the link between TAM and diabetes is strong, metabolic effects of aromatase inhibition are less consistent ^56,57^, potentially because these drugs are relatively new and studies have not been adequately powered or compared this treatment to untreated women. Aromatase inhibitor treatment was recently reported to elevate risk for diabetes ^10^, insulin resistance, and fat gain ^58^, and to increase non-alcoholic hepatic steatosis regardless of BMI ^41^. Our observation of decreased adipocyte progenitor cells and subsequent adipocyte hypertrophy is consistent with a long-term risk of type 2 diabetes due to inappropriate lipid storage. In the context of chronic positive energy balance, a shift towards adipocyte hypertrophy would be the only means of adipose tissue expansion, which is consistent with data from our mouse model as well as the adipose tissue from women treated with TAM. Although it may seem counterintuitive, the formation of new adipocytes in expanding adipose tissue maintains the metabolic health of the organism by preventing metabolic derangements associated with ectopic lipid deposition ^59,60^. Importantly, human studies are limited to those in women who were diagnosed with cancer, where obesity is a confounding risk factor for both diabetes and breast cancer. We show in a preclinical model that breast cancer endocrine therapy, in the absence of cancer, promotes dysregulated metabolism particularly in HFHS fed females with excess adiposity.

*Esr1* was highly expressed in mesenchymal stem cells and a myeloid-derived progenitor cell population from adipose tissue (Figure 3). Others have shown that *Esr1* expression was elevated in primary FACS-isolated mouse adipose progenitor cells compared to either the total stromal/vascular fraction (made up largely of preadipocytes) or to mature adipocytes ^61^. We found that endocrine therapy appeared to stimulate early steps of *de novo* adipocyte differentiation in subcutaneous adipose tissue regardless of diet; but, in HFHS fed females, the progenitor cells that give rise to preadipocytes did not increase. Consistent with our observations, TAM was reported to inhibit proliferation and subsequent adipogenic differentiation of primary human adipose progenitor cells in vitro ^62^. Notably, complete ablation of *Esr1* from adiponectin-positive mature adipocytes also promoted increased adiposity ^63^. Together, data from our study and others support a prominent role for ER signaling in maintaining adipose progenitors and suggest potentially fundamental differences in progenitor cell self-renewal capabilities between females with and without obesity. As predicted based on the depletion of adipocyte progenitors, by seven weeks of treatment adipocyte size was greater with TAM and EWD compared to E_2_ in the HFHS mice, which was accompanied by hepatic steatosis and elevated HOMA-IR. Whether these effects persist after discontinuation of TAM, which would resemble a clinically relevant scenario, is the focus of future studies.

To model aromatase inhibition, we withdrew supplemental estradiol (EWD) from OVX females, lowering circulating estrogen levels substantially based on our previous in-depth analysis of adipose tissue *Cyp19a1* expression and E_2_ in mice and rats ^26,27^. The ability of rodent adipose to aromatize androgens to estrogens remains controversial in the fields of obesity and cancer ^64,65^. Using the EWD approach, we observed metabolic outcomes consistent with those reported in breast cancer patients ^58^. EWD had similar effects in LFLS and HFHS females: greater fat accumulation, food intake, and hepatic lipid deposition; however, despite gaining body fat, LFLS-fed females maintained fasting insulin and glucose levels similar to E_2_ treated females. These data highlight that weight gain is not always accompanied by deregulated metabolism when the adipocyte progenitor population is maintained, and adipocyte size remains small. As we expected based on the depletion of adipocyte progenitors, after seven weeks of endocrine therapy, HFHS females suffered the worst consequences with excess fat gain, hepatic steatosis, and impaired glucose tolerance. Whether the effects of cancer therapy on the liver or whole-body insulin sensitivity are due to lipid spillover from hypertrophic adipocytes in mice or in humans is unclear.

One important aspect of our study was that we investigated the metabolic effects of TAM using a relatively low dose, known to effectively inhibit the growth of endocrine-sensitive ER-positive breast tumors ^24,25^. A limitation of our approach is the difficulty in precisely modeling the specific ratio of estrogens to tamoxifen as they would occur in women throughout their menstrual cycles. In addition, we delivered tamoxifen in a subcutaneous pellet; however, this has been shown to result in effective inhibition of human ER-positive breast tumor growth ^24,25^. Women are prescribed 20-40 mg TAM per day, as either a single or divided dose, delivered orally. Assuming the average adult female mass is approximately 76 kg, this results in a dose of 0.26-0.52 mg/kg/day. We administered TAM as a subcutaneous 5 mg pellet, released over 60 days. For the mice in our study, this results in an average daily dose of 3.0-3.6 mg/kg/day, based on body mass range. While that daily dose of TAM exceeds what is given to breast cancer patients, we did not observe weight loss, which could indicate acute toxicity. Previous studies in rodents have used doses of TAM designed to activate expression of Cre-ER transgenes, ranging from 25-300 mg/kg/day administered over a few days by IP injection or oral gavage ^19-22,66-70^, and many have been done in males. Major metabolic outcomes reported in published studies include decreased food intake ^66^, rapid adipose tissue loss ^19-21^, adipose tissue browning ^21,32^, and hepatic steatosis ^68,69^. With the exception of one study conducted at cold temperature ^32^ and one conducted on high fat fed mice ^21^, most were done on chow-fed males at ambient temperature (20-24C). In contrast, we performed studies on mice at thermoneutrality (∼30C) and compared HFHS-to LFLS-fed females. Overall, our findings of adipose accumulation and inappropriate adipose tissue expansion are consistent with a later life risk for type 2 diabetes.

A second objective of this study was to use our preclinical model to determine if interventions that target insulin action could be effective during endocrine treatment to prevent the metabolic disturbances associated with endocrine therapy. To this end, we administered one of two widely used anti-hyperglycemic drugs, metformin or pioglitazone, each of which had distinct effects depending on the endocrine therapy and the outcome measure (summarized in Supplemental Table 2). Females treated with TAM appeared to benefit from either intervention, with main differences seen in hepatic steatosis. During EWD, these same benefits were not seen, suggesting that the hormonal environment may influence the response of tissues to drugs that improve insulin action. In mouse 3T3-L1 preadipocytes, E_2_ treatment increased levels of PPARγ protein. The combination of E_2_ and pioglitazone also increased PPARγ levels compared to pioglitazone treatment alone ^71^; however, E_2_ was also shown to inhibit the pro-adipogenic effects of troglitazone on 3T3-L1 cells, albeit at a high concentration (100 µM) ^72^. PPAR and ER family members can bind the same DNA response elements ^73^ but it is unclear how PPARγ signaling involves ERα in adipocyte progenitors, and whether potential crosstalk is modulated by estrogens or tamoxifen. Potential sex differences in the response to anti-hyperglycemic drugs were reported for some patients with type 2 diabetes, where women benefitted more from thiazolidinedione treatment than men while the inverse was seen with sulfonylureas ^74^. In general, data are limited on sex- or menopausal status-specific benefits of anti-hyperglycemic therapies with any outcome measure. Our study suggests the need for careful patient selection, potentially considering menopausal status and currently prescribed cancer therapy when implementing metabolic interventions.

Overall, E_2_-activated ER signaling is known to impact a variety of tissues and cell types at different stages of differentiation. We posit that one important role is to maintain the adipocyte progenitor pool (Figure 7), which allows healthy adipose tissue expansion by hyperplasia. In the context of obesity when progenitors are depleted and ER signaling is disrupted either by withdrawal of the E_2_ ligand or in the presence of an antagonist, adipogenesis occurs without repopulation of preadipocytes. Over time, especially in the presence of a chronic positive energy balance, hypertrophic adipocytes reach their limit of storage capabilities, which can result in ectopic lipid deposition. Anti-estrogen therapies have been instrumental in preventing recurrence of breast cancer for the vast majority of patients. With increased survival of breast cancer patients and the heightened awareness of elevated type 2 diabetes risk in this population, our study suggests the need for close monitoring and potential anti-hyperglycemic intervention for some women during endocrine therapy.

**Figure 7.**
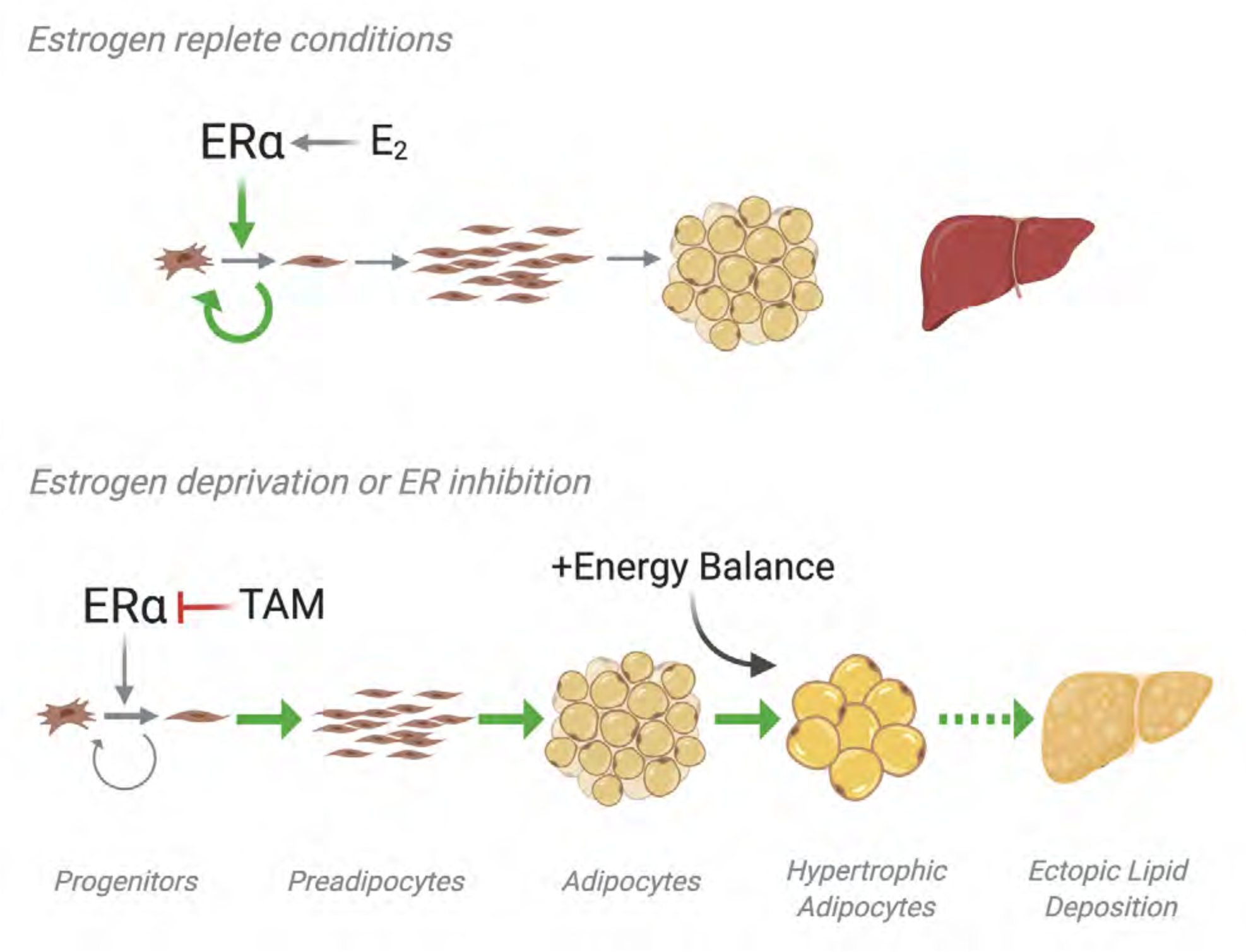
Proposed model of endocrine therapy effects on adipose tissue. ERα signaling maintains adipocyte progenitor pools and inhibits preadipocyte expansion, adipocyte differentiation, and hypertrophy. Disruption of E_2_ signaling through either tamoxifen treatment or withdrawal of E_2_ depletes the adipocyte progenitor pool causing adipocyte hypertrophy consistent with a phenotype that precedes insulin resistance and the development of type 2 diabetes.

## Methods

### Bioinformatics

Single cell RNA sequencing data were accessed from Tabula Muris (https://tabula-muris.ds.czbiohub.org/) in April 2020. Expression of *Esr1, Esr2*, and *Cyp19a1* was examined and visualized in FACS processed cells from all tissues together, and from adipose (fat), limb muscle, liver, and mammary gland individually. Plots were exported from the website, and numeric data were used to generate a heatmap for *Esr1* and *Esr2* expression in Graph Pad Prism 8.

### Mice

Female C57BL/6J mice were purchased from Jackson Laboratories at 6 weeks of age (stock #000664). Mice acclimated for 1 week and were then given either low fat/no sucrose diet (LFLS; Research Diets D11092101) or high fat (40%)/high sucrose (292.5g/3902kCal) diet (HFHS; Research Diets D15031601) *ad libitum*. After 2 weeks in standard ventilated housing, mice were moved to warm water blankets set at 42C, to maintain an internal cage temperature of ∼30C, as we have previously described ^26^. At 14 weeks of age, mature mice were ovariectomized under isoflurane anesthesia and immediately supplemented with 17β-estradiol (E_2_), provided in the drinking water at a final concentration of 0.5 µM. E_2_ supplementation continued for 2 weeks, at which time mice were randomized based on body fat percentage by qMR Echo (ECHO MRI) within diet groups to one of 3 treatments: E_2_-maintenance (E_2_), E_2_ plus tamoxifen (E_2_+TAM), or E_2_ withdrawal (EWD). Tamoxifen free base was delivered by subcutaneous pellet implant, with each mouse receiving a 5 mg, 60-day release pellet (Innovative Research of America). Mice were treated for either two or seven weeks, then fasted for 4-6 hours and euthanized according to approved AAALAC guidelines. In a separate study, HFHS fed mice were matured as described, and then randomized based on body fat percentage to receive E_2_+TAM or EWD, and one of two interventions during treatment. Metformin was delivered for 7 weeks in the drinking water at 2 mg/mL, as we have done previously ^75-77^. Pioglitazone was administered in the diet (Research Diets D15301601) at 0.1 grams/kg diet, as described ^78^. At the end of study, blood and tissues were collected, and organs were weighed. All animal studies were approved by the University of Colorado Denver Institutional Animal Care and Use Committee.

### Whole-animal Calorimetry

Whole-body calorimetry was performed on a subset of LFLS- and HFHS-fed females, during the second week after initiation of endocrine therapy. Mice were acclimated for 3 days and data were collected for an additional 24 hours. Total energy expenditure, resting energy expenditure and spontaneous physical activity were measured in a metabolic monitoring system (Oxymax CLAMS-8M; Columbus Instruments, Columbus, OH). Individual metabolic cages included an animal activity meter (Opto-Max, Columbus Instruments, Columbus, Ohio) to allow for the calculation of total, ambulatory and non-ambulatory activity by monitoring beam breaks within a one-dimensional series of infrared beams. Metabolic rate was calculated from gas exchange measurements acquired every 18 minutes using the Weir (1949) equation: MR = 3.941 x VO_2_ + 1.106 x VCO_2_ – 2.17 x N, where N is urinary nitrogen (61). MR was averaged and extrapolated over 24 hours to estimate total energy expenditure. Energy intake was measured daily while the mouse was in the metabolic monitoring system. Energy balance was calculated as the difference between energy intake and expenditure (n=4 per group).

### Mouse tissue analysis

Tissues were fixed in 10% neutral buffered formalin, then processed and embedded using standard histology procedures. Five µm sections were stained with H&E to visualize tissue morphology and for quantification of adipocyte size distribution, which was performed using the Adiposoft plugin ^79^ and ImageJ software. For each group, one section from at least 3 mice (500-1800 cells per mouse) was analyzed. Liver sections were stained with H&E and hepatic steatosis scores (0-3) were assigned by a Board-Certified Liver Pathologist, blinded to experimental groups. For each group, sections from 3-7 mice were evaluated. Pancreas sections were stained using a guinea pig polyclonal anti-insulin antibody (Abcam ab7842) and counterstained with hematoxylin. Stained sections were scanned and analyzed using the Aperio Digital Pathology system (Leica Biosystems) and the positive pixel count algorithm quantified percent positive of total pixels in full representative sections.

### Human tissue analysis

Breast adipose tissue was collected by the Komen Tissue Bank and the IU Simon Cancer Center (https://virtualtissuebank.iu.edu/). Tissues were formalin fixed, processed, embedded, cut, and stained with H&E. Sections were scanned using the Aperio Digital Pathology system (Leica Biosystems). Images were accessed in May 2020 through the Aperio Digital Pathology system (Leica Biosystems) and adipocyte size was quantified using the Adiposoft plugin ^79^ and ImageJ software. Between 350 and 2000 cells were analyzed per tissue section.

### Mitochondrial Respiration

Mitochondrial respiration was measured using Oroboros Oxygraph-2k (O2k, OROBOROS INSTRUMENTS Corp., Innsbruck, Austria) according to modifications from previously described protocols ^80-82^. Immediately after tissue harvest, ∼100 mg of liver tissue (left lobe) was placed in ice cold mitochondrial preservation buffer [BIOPS (10 mM Ca-EGTA, 0.1 mM free calcium, 20 mM imidazole, 20 mM taurine, 50 mM K-MES, 0.5 mM DTT, 6.56 mM MgCl2, 5.77 mM ATP, 15 mM phosphocreatine, pH 7.1)] and kept on ice. Pieces of liver tissue (2-3 mm x 2-3 mm) were removed and separated mechanically (in BIOPS and on ice) and partially teased apart by fine forceps (Dumont 5, non-magnetic). Liver tissue was blotted dry on Whatman paper for a wet weight measurement (∼1 mg) and added to mitochondrial respiration buffer [MiR05 (0.5 mM EGTA, 3 mM magnesium chloride, 60 mM K-lactobionate, 20 mM taurine, 10 mM potassium phosphate, 20 mM HEPES, 110 mM sucrose, 1 g/L bovine serum albumin, pH 7.1)] that had been prewarmed to 37°C in the chamber of the O2k. The liver tissue was permeabilized in the O2k chamber by incubation with digitonin (4 µg/mL for 15-20 min). Oxygen concentration in the MiR05 was started at approximately 400 µM and runs were completed before oxygen levels dropped below 220 µM

Two sets of substrates and inhibitors were added to assess respiration rates at several states (Run 1 and 2). Rates for Run 1 were measured following the addition of 5 mM pyruvate (P) and 1 mM malate (M) (state 2 PM); PM with 2 mM adenosine diphosphate (ADP) (state 3 PM); PM, ADP and 10 mM glutamate and 10 mM succinate (S) (state 3 PMGS); PMGS, ADP and 2 µg/ml oligomycin (state 4 PMGS). Rates for Run 2 were measured following the addition of 50 μM palmitoylcarnitine (PC) and 1 mM malate (M) (state 2 PCM); PCM with 2 mM adenosine diphosphate (ADP) (state 3 PCM); PCM, ADP and 10 mM succinate (state 3 PCMS); PCMS, ADP and 2 µg/ml oligomycin (state 4 PCMS). Uncoupling was performed at the end of each run by 1-2 uM stepwise titration of FCCP (carbonyl cyanide 4-(trifluoromethoxy)phenylhydrazone). Respiration control ratios (RCR) were calculated as a ratio of state 3 PM/state 2 PM (RCRPM), state 3 PMGS/state 4 PMGS (RCRPMGS), state 3 PCM/state 2 PCM (RCRPCM), state 3 PCMS/state 4 PCMS (RCRPCMS).

### Western Blotting

Liver (35–50mg) was flash frozen and stored at −80°C until it was homogenized 1:20 in mammalian lysis buffer (MPER with 150 mM sodium chloride, 1 mM of EDTA, 1 mM EGTA, 5 mM sodium pyrophosphate, 1 mM sodium orthovanadate, 20 mM sodium fluoride, 500 nM okadaic acid, 1% protease inhibitor cocktail). After homogenization, samples were centrifuged at 18,000g for 10 minutes at 4°C. The resulting supernatant was analyzed for protein concentration by Bradford protein assay. Protein samples (10 µg to 25 µg) in Laemmli sample buffer were run on 4-20% Tris-HCl gels. The resolved proteins were electrophoretically transferred to PVDF membranes, and equivalence of protein loading was assessed by staining of membrane-bound proteins by Ponceau S stain. Blots were probed using an antibody against one subunit of each of the mitochondrial oxidative phosphorylation complexes (Abcam #ab110412; 1:1000; overnight at 4°C) and followed by fluorescent secondary (1:10,000, 1 hr at room temperature). Proteins were detected by Li-COR (Odyssey CLX) Western blot scanner, and densitometric analysis was performed using Image Studio v4.1.

### Adipose Tissue FACS

Whole inguinal subcutaneous adipose depots were excised, minced briefly (5 min) with dissecting scissors, and digested for 75 mins at 37C in a collagenase solution (Collagenase type II, Worthington LS004177). (HBSS with 3% BSA, 0.8mM ZnCl2, Mg, Ca, 0.8mg/ml collagenase). Stromal pellets were incubated in red blood cell lysis buffer (Sigma Aldrich), washed, and stained with fluorescent-conjugated antibodies for CD45, CD31, Sca1, CD29, CD34, and CD24 as described ^40^. A full list of antibody dilutions, sources, and fluorophores is available in Supplemental Table 3.

### Serum Analyses

One week prior to sacrifice, oral glucose tolerance tests were performed on a subset of mice from each group. Mice were fasted for 4 hours and then given 5 g/kg body mass dextrose from a 50% dextrose solution (VetOne, Boise, ID) by oral gavage. Blood glucose levels were measured using glucometers (Contour, Parsippany, NJ) with tail vein blood drawn at 0, 10, 20, 40, 60, 80, and 120 minutes post treatment.

Serum was prepared from fasted blood collected at end of study. Insulin (Alpco Mouse Insulin ELISA; 80-INSMS-E01) and glucose (Cayman Chemicals Glucose Colorimetric Assay; 10009582) were measured in technical duplicates according to manufacturers’ protocols. The homeostatic model assessment of insulin resistance (HOMA-IR) was calculated according to the formula: fasting insulin (µU/L) x fasting glucose (mmol/L)/22.5.

### Statistics

The purpose of this study was to determine the impact of endocrine therapies on metabolic health in the context of obesity. To address this objective, the statistical analyses were designed a priori to compare each endocrine therapy to the E_2_ alone condition, as that provides the most clinically relevant assessment of the data. Therefore, two separate two-way analysis of variance (ANOVA) were used. The first ANOVA assessed the effect of diet (LFLS vs. HFHS) and EWD treatment (EWD vs. E_2_). The second ANOVA assessed the effect of diet (LFLS vs. HFHS) and TAM treatment (E_2_ vs. E_2_+TAM). Pairwise comparisons were performed using the Tukey test where appropriate. For metformin and pioglitazone interventions, data were compared between intervention and control mice within each endocrine therapy group using unpaired t-tests. The level of statistical significance was set at *P* < 0.05. Data are expressed as mean ± SE.

### Study Approval

All mouse studies were approved by the University of Colorado Institutional Animal Care and Use Committee. Collection of human breast adipose tissue was approved by the Indiana University Institutional Review Board for use in the Komen Tissue Bank.

## Author Contributions

RLS and EAW designed and oversaw the study and collected and analyzed data from mouse and human experiments. RMF, SEH, LAK, SJM, FK, BF, GV, JAH, GJ, AMYZ collected, assembled, and interpreted data from mouse experiments. JK evaluated and scored liver samples. JDJ, PSM, JEBR, RLS and EAW interpreted data and wrote and edited manuscript. SWH inspired the study from the patient perspective and provided input throughout the project and on the manuscript.

## Acknowledgements

Data from the Susan G. Komen Tissue Bank at the IU Simon Cancer Center was used in this study. We thank contributors, including Indiana University who collected data used in this study, as well as donors and their families, whose help and participation made this work possible. We also thank Veronica Wessels for providing outstanding histological expertise.

