## Supplemental Figures and Tables for "Breast cancer endocrine therapy exhausts adipocyte progenitors promoting weight gain and glucose intolerance"

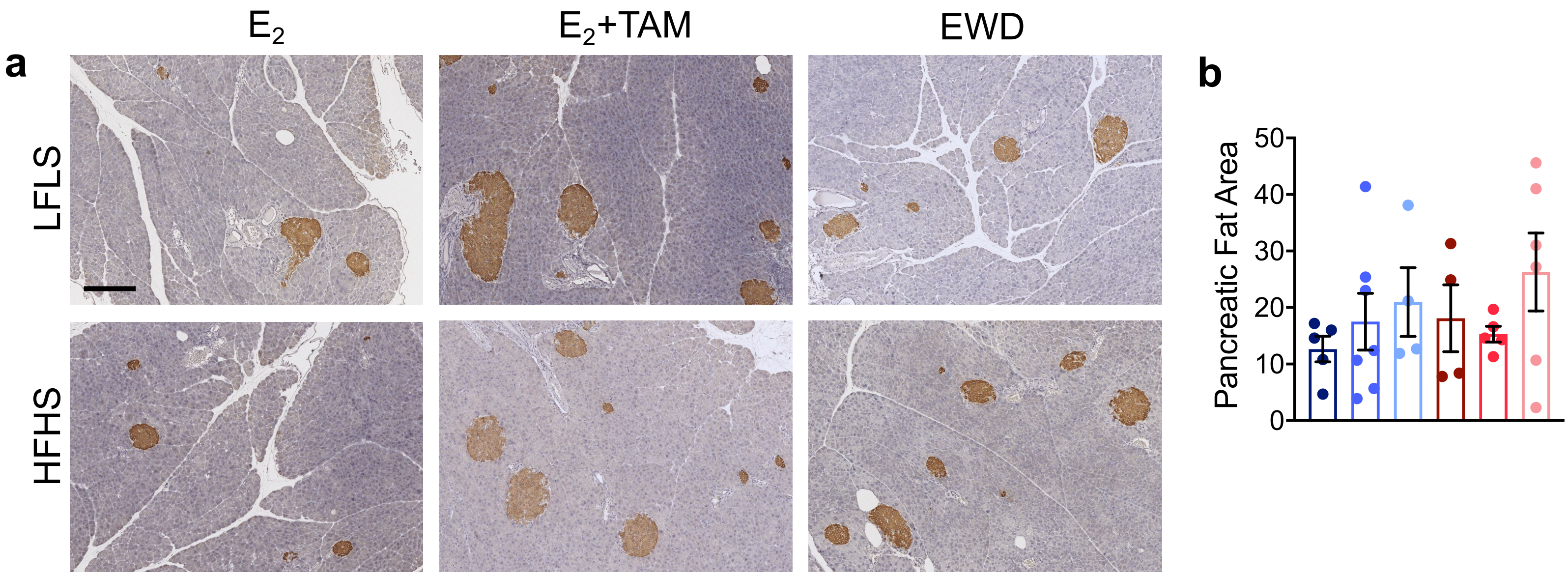


**Supplemental Figure 1. Endocrine therapy associates with increased insulin-positive cells in the pancreas. a**, images of pancreas from one animal in each group showing insulin positive cells measured by IHC. **b**, pancreas fat area quantified in sections from LFLS $E_2$ (n=5), LFLS $E_2$+TAM (n=6), LFLS EWD (n=4), HFHS $E_2$ (n=4), HFHS $E_2$+TAM (n=6), HFHS EWD (n=6).

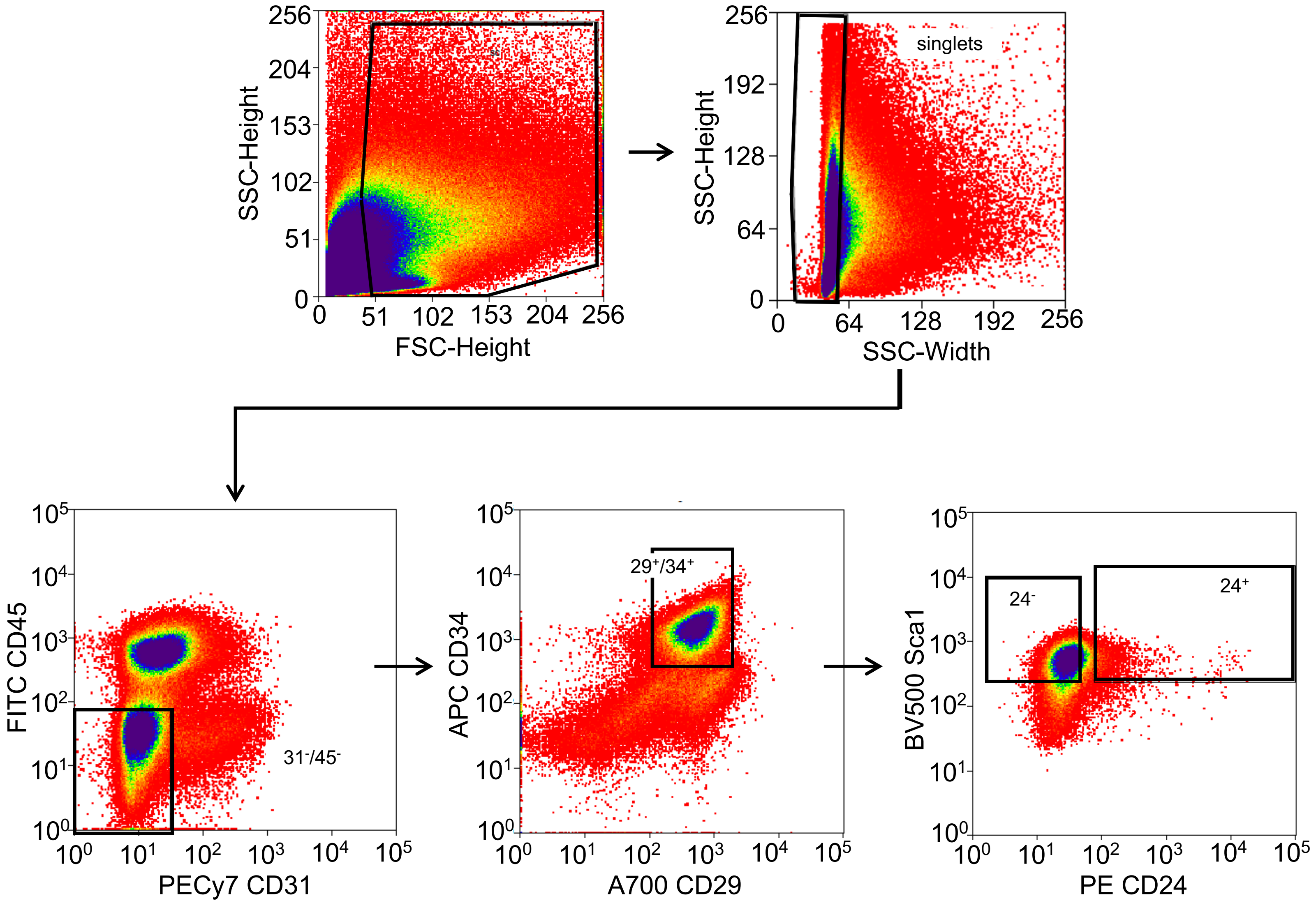


**Supplemental Figure 2. Representative FACS plots for preadipocytes and progenitors.** Flow sorted cells for each animal were collected with gates set using appropriate fluorescence minus one and unstained controls.

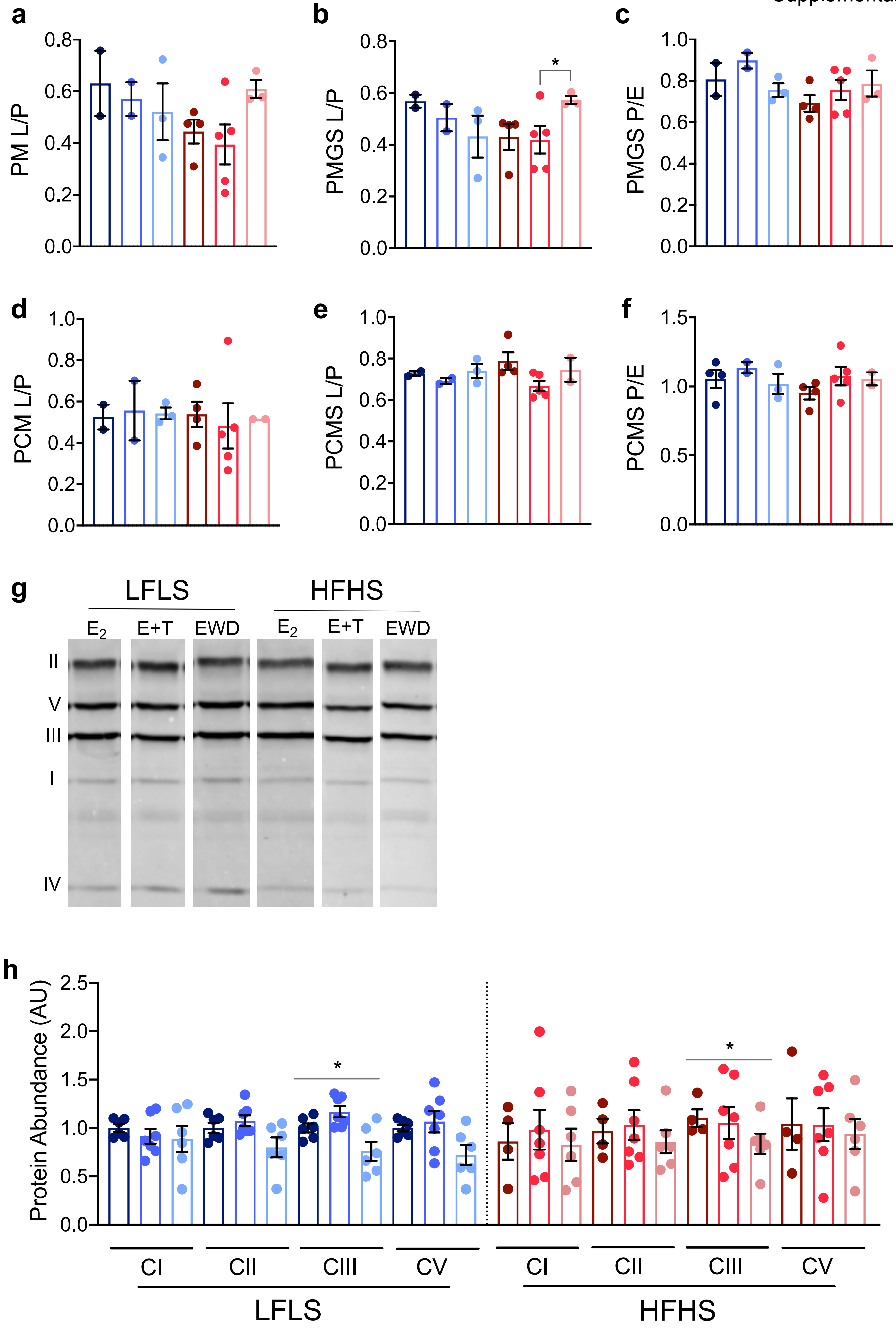
a
PM L/P
0.0
0.2
0.4
0.6
0.8
b
PMGS L/P
*
c
PMGS P/E
1.0
d
PCM L/P
e
PCMS L/P
f
PCMS P/E
0.5
1.5
g
LFLS
HFHS
E2
E+T
EWD
II
V
III
I
IV
h
Protein Abundance (AU)
2.0
2.5
CI
CII
CIII
CV
LFLS
HFHS

**Supplemental Figure 3. Endocrine therapy does not influence hepatic mitochondrial content or function.** Mitochondrial respiration was assessed with both a carbohydrate-linked SUIT (Palmitate: P, Malate: M, Glutamate: G, and Succinate: S) and a lipid-linked SUIT (Palmitoylcarnitine: PC, M, and S). a, ratio of leak to ADP stimulated respiration (L/P ratio) with PM. b, L/P ratio with PMGS. This ratio was greater in the HFHS mice with EWD (*P* = 0.04). c, the ratio of ADP stimulated respiration to uncoupled respiration (P/E ratio). d-f, respiration ratios with PCMS suit. g, representative immunoblot of one mouse per group. h, quantification of mitochondrial content measured by western blot. Data are presented as AU with representative blot. Complex III was lower with EWD (P =0.012). No other effects of diet or treatment were observed in other mitochondrial complexes.

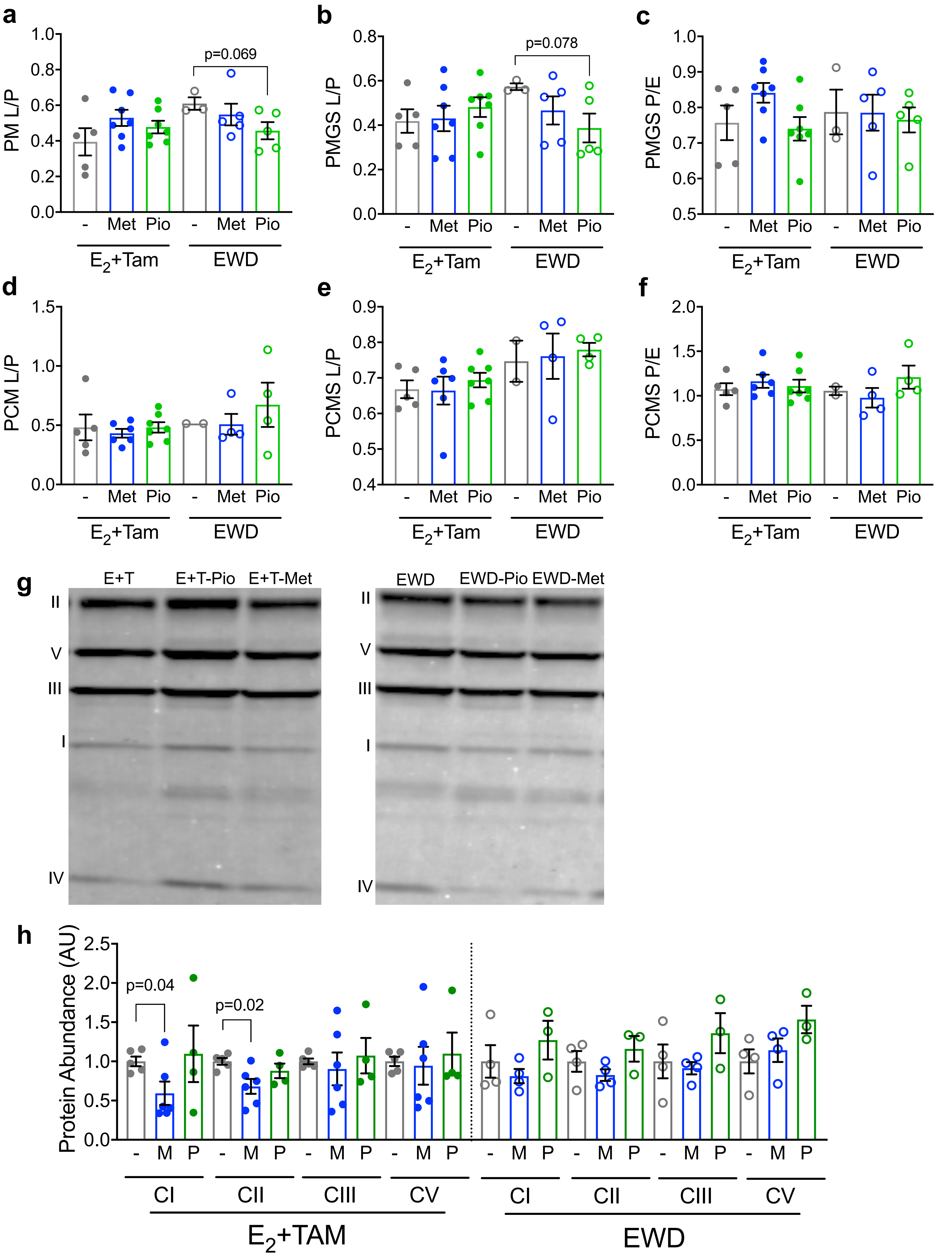
a
PM L/P
p=0.069
b
PMGS L/P
p=0.078
c
PMGS P/E
d
PCM L/P
e
PCMS L/P
f
PCMS P/E
- Met Pio - Met Pio
$E_2$+Tam
EWD
g
E+T
E+T-Pio
E+T-Met
EWD
EWD-Pio
EWD-Met
II
V
III
I
IV
h
Protein Abundance (AU)
p=0.04
p=0.02
- M P
CI
CII
CIII
CV
$E_2$+TAM
EWD

**Supplemental Figure 4. Anti-hyperglycemic interventions do not impact hepatic mitochondrial content or function.** Mitochondrial respiration was assessed with both a carbohydrate-linked SUIT (Palmitate: P, Malate: M, Glutamate: G, and Succinate: S) and a lipid-linked SUIT (Palmitoylcarnitine: PC, M, and S). **a**, ratio of leak to ADP stimulated respiration (L/P ratio) with PM. **b**, L/P ratio with PMGS. **c**, the ratio of ADP stimulated respiration to uncoupled respiration (P/E ratio). **d-f**, respiration ratios with PCMS suit. **g**, Mitochondrial content measured by western blot. Data are presented as AU with representative blot. Complex I was lower with metformin treatment in the context of TAM (P =0.04). No other effects of diet or treatment were observed in other mitochondrial complexes.

**Supplemental Table 1.** Summary of metabolic measures in LFLS and HFHS fed females treated with $E_2$+TAM or EWD.

| | LFLS | | HFHS | |
|---|---|---|---|---|
| ***Outcome*** | $E_2$+TAM | EWD | $E_2$+TAM | EWD |
| **Body Fat (mg/g)** | - | ↑ | ↑ | ↑ |
| **HOMA-IR** | - | - | ↑ | ↑ |
| **Adipocyte Size** | - | ↑ | ↑ | ↑ |
| **Preadipocytes** | ↑ | ↑ | ↑ | ↑ |
| **Adipocyte Progenitors** | - | ↑ | ↓ | - |
| **Hepatic Steatosis** | - | ↑ | - | ↑ |

**Supplemental Table 2.** Summary of metabolic measures in HFHS fed females treated with $E_2$+TAM or EWD, and with either metformin (Met) or pioglitazone (Pio).

| | **$E_2$+Tamoxifen** | | | | **EWD** | | | |
|---|---|---|---|---|---|---|---|---|
| | Fat Mass (g) | HOMA-IR | Adipocyte Diam (µm) | Hepatic Steat | Fat Mass (g) | HOMA-IR | Adipocyte Diam (µm) | Hepatic Steat |
| **Met** | **-** | ↓ | ↓ | **-** | **-** | **-** | **-** | **-** |
| **Pio** | ↓ | ↓ | ↓ | ↓ | **-** | ↑ | ↓ | **-** |

**Supplemental Table 3.** Antibodies for FACS analysis of adipose tissue.

| Antibody | Color | Dilution | Vendor/Cat # |
|---|---|---|---|
| CD29 | AF 700 | 1:200 | Biolegend 102218 |
| CD31 | PE Cy7 | 1:500 | Biolegend 102418 |
| Sca-1 | V500 | 1:500 | BD 561229 |
| CD45 | FITC | 1:1000 | Biolegend 103108 |
| CD24 | PE | 1:100 | Biolegend 138504 |
| CD34 | APC | 1:50 (2 in 100ul) | Biolegend 119310 |
